# Between Behaviors: Comparison of Two Dynamical Models of Behavioral Switching for *C. Elegans* Locomotion

**DOI:** 10.64898/2026.02.26.708303

**Authors:** Denizhan Pak, Randall D. Beer

## Abstract

Organisms must manage a trade-off between robustness and flexibility as they enact adaptive behaviors. One way organisms achieve this is by navigating a network of quasi-stable behavioral states. Evidence for such behavioral states has been observed in many organisms, and new methods for detecting these states have taken on a prominent research focus. Although dynamical models demonstrating behavioral switching have been developed significantly over the past few decades, theories of the similarities and differences among these models, necessary for advancing empirical modeling, have not yet been fully elaborated. Here, we consider behavioral switching in two different classes of dynamical models of the forward-reversal behavioral transition in *C. elegans*. We first show how fundamentally different models can give rise to similar phenomena under noisy conditions. We then analyze the deterministic aspects of these models to expand on their differences, clarifying the theoretical relationship between them. Finally, we demonstrate how sequence models can be further extended to incorporate dwell times for behavioral states. Our work contributes toward a broader theoretical understanding of behavioral switching in adaptive systems.

## 1 Introduction

Organisms must perform adaptive behaviors to maintain conditions conducive to their survival in constantly changing environments. These behaviors emerge from the interaction of heterogeneous systems operating across multiple spatial and temporal scales, often exhibiting long-lived complex nonlinear transient dynamics [1]. But behavior remains remarkably structured, with organisms utilizing a limited range of temporary yet repeated patterns of movement that are sequenced together. This organization suggests that behavior is composed through recombinations of a more restricted set of underlying primitives [2].

Substantial evidence for such primitives comes from advances in computational ethology. In particular, pose-tracking algorithms [3, 4] have greatly expanded our ability to extract sequences of primitives directly from video recordings [5]. At the neural level, specific neural ensembles have been found to activate in sequences that align with the corresponding behavioral primitives [6], further supporting the view that complex behavior is structured from reusable dynamical building blocks.

Although significant advances in empirical research have been made, our theoretical understanding of behavioral sequences remains limited. Behavioral sequences are often modeled using Markov chains, in which individual primitives are treated as discrete states with probabilistic, instantaneous transitions between them [5, 7, 8]. Markov models imply a stationary distribution over behavioral primitives, which is inconsistent with experimental evidence—even in simple organisms such as *C. elegans*, which exhibit context-dependent state distributions [9]. One practical solution is to use Markov chains operating on timescales much shorter than the behavioral sequences of interest. However, even these models fail to capture non-Markovian properties revealed at finer scales [7].

This limitation reflects a deeper problem with describing behavior purely in terms of discrete states [10]. An alternative is to describe behavior as a continuous trajectory through the phase space of the brain–body–environment system [11]. Ordinary differential equations (ODEs) provide a natural framework for such a description. In this view, primitives correspond to specific subregions in phase space which are revisited. The fundamental units of behavior thus become recurring patterns which emerge from underlying biophysical mechanisms rather than fixed modeling choices. However, explaining the emergence of recurrent subregions requires additional mathematical machinery [12].

The region of phase space corresponding to a behavioral primitive must be stable enough to ensure repetition and robustness, yet unstable enough to permit escape and switching; following [13], we consider these regions to be quasi-stable. Recent work has identified several dynamical mechanisms capable of producing quasi-stable behavior [12–14].

Quasi-stability can arise from a multistable system interacting with external energy and noise, where perturbations drive sequential dynamics. However, defining behavior primitives strictly as stable attractors neglects spontaneous activity and internal noise, both central to biological systems [12]. Alternatively, metastability has been proposed as an organizing principle for quasi-stability. Topologically, metastability may emerge through generalized saddle structures (e.g., homoclinic or heteroclinic channels) or ghost states [15]. These mechanisms differ in structural requirements and analytical tractability, yet both can generate metastable dynamics. Additional complexity arises in phenomena such as chaotic itinerancy or Milnor attractors, where these mechanisms interact to produce more complex, potentially chaotic dynamics [14].

Although these mechanisms have been individually characterized and applied to brain and behavior, they have not been directly compared in explaining the same behavioral phenomenon. A robust theory of behavioral switching requires principled distinctions and clearly defined ranges of applicability for these mathematical frameworks. Here, we compare mechanisms of metastability in two dynamical models of behavioral switching within a biologically relevant context.

## 2 A Motivating Example: *C. Elegans*

*C. elegans* offers a relatively simple biological example that will allow us to clarify the theoretical complexities of behavioral sequences. The nematode is one of the few organisms for which a nearly “complete” connectome has been identified [17]. The hermaphroditic variant, often used in scientific research, has only 302 neurons [18] and has been studied thoroughly as a model organism [19]. The experimental tractability and existing knowledge surrounding the nematode make it an ideal model system for identifying theoretical complexities that will be faced as our models and theories scale to more complex organisms.

During locomotion, *C. elegans* exhibits distinct wave patterns across its body that allow it to move in a viscous environment [20]. Each of these wave patterns corresponds to a different behavioral state [21]. Although the exact number and types of modes have been under debate, a consistent finding is the occurrence and switching between two different modes: forward and reverse locomotion. As *C. elegans* navigates, its primary forward motion is punctuated by reversals in which it moves backwards for short bursts of time.

The neural basis of this phenomenon has become a significant topic of interest for several reasons. Firstly, the global neural dynamics of *C. elegans* seem to cycle through a small network of modes that correspond to its behavioral states [6], two of which are forward and reversal. Secondly, the forward and reversal behaviors require overlapping neuromechanical circuitry, which poses an interesting question about the relationship between structure and function [22]. Finally, it has been shown that by modifying the frequency and duration of these behavioral states, the worm is able to implement different foraging search strategies [23].

Various models have been proposed to explain this behavioral sequence [24], both at the neural [8] and neuromechanical levels [25]. These models differ in their level of abstraction and biological detail. Roberts et al. in [8], proposed that the forward and reversal behavioral states correspond to excitation of two different clusters of neurons: one consisting of AVBL, AVBR, PVCL, and PVCR neurons correlated with forward movement, and the other AVAL, AVAR, AVDL, AVDR, AVEL, and AVER cluster linked to reversals (Figure 1). They also proposed a quantitative Markov model that was able to match empirical observations at the behavioral and neural levels (Figure 2).

**Fig. 1.**
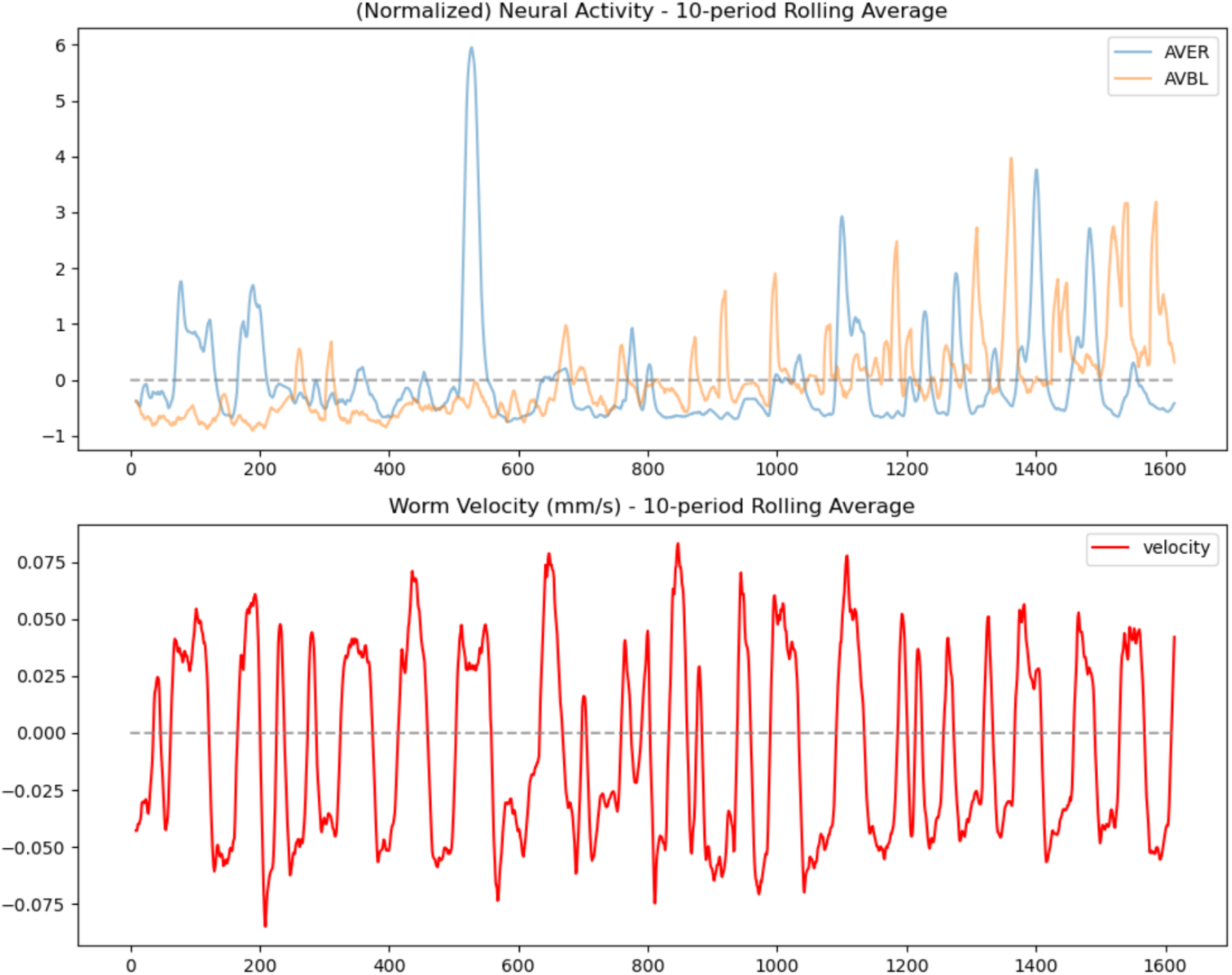
Neural dynamics of a representative neuron from the forward cluster (AVER) and neural dynamics of a neuron from the reversal cluster (AVBL) as well as the velocity of the freely moving *C. elegans*. The velocity of the worm is tightly correlated with neural activity from neurons of the corresponding cluster. The data plotted is adapted from the WormWideWeb project [16].

**Fig. 2.**
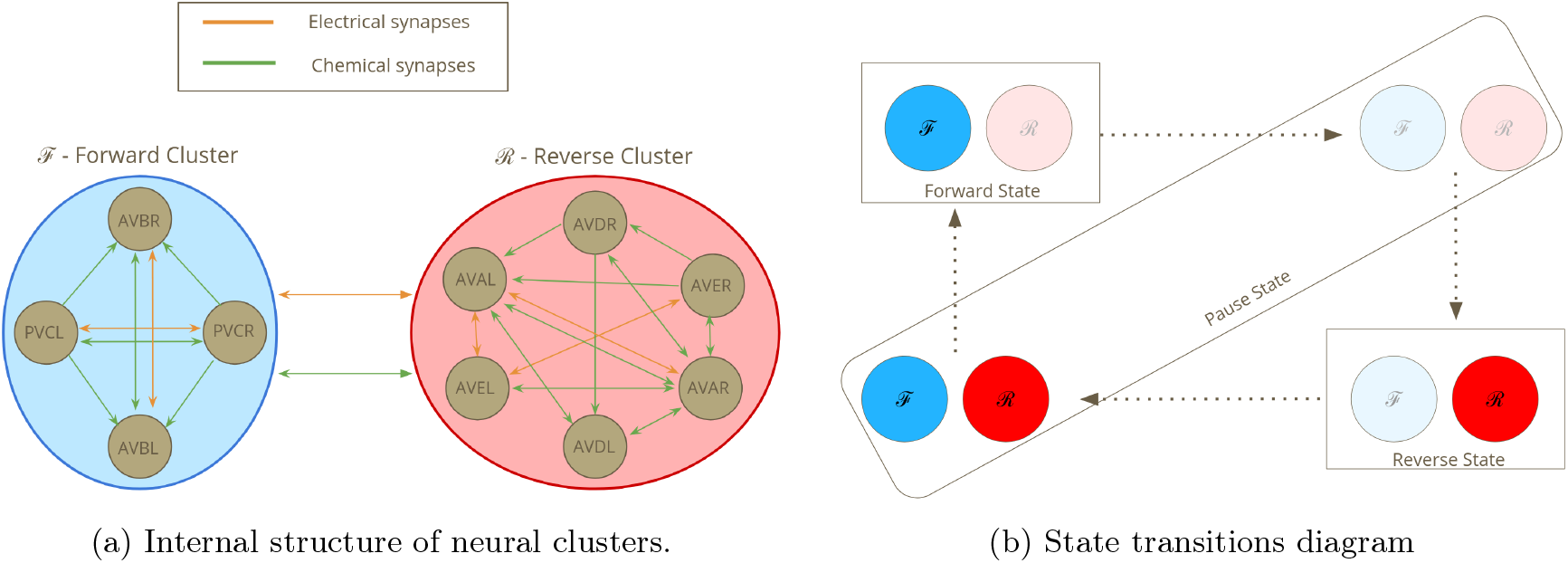
The conceptual model underlying forward-reversal transitions as proposed in [8]. The model depends on two clusters of neurons: one for forward movement (ℱ) and one for reversals (ℛ). The activity of these clusters results in three different behavioral states: forward, reversal, and pauses. Pauses occur during transitions between forward and reversal states. Although behaviorally three states are observed, the model suggests that the same behavioral state of pause can be caused by two different neural states: where both clusters are active (transitioning from reversal to forward) or both clusters are inactive (transitioning from forward to reversal).

To explain the behavioral data, they found that the most appropriate model consisted of three-sates: forward, pause, and reversal. However, they also found that the best model to explain the neural data was a four-state rather than three-state model. While the forward and reversal behavioral states could be understood as the excitation of the corresponding clusters, they suggested that the pause states occur either due to excitation in both clusters in the forward-to-reversal transition or due to inactivity of both clusters in the reversal-to-forward transition. This approach allowed for a three-state description of the behavior and a four-state description of the neural activity.

The model of Roberts et al. was a good fit for the empirical data and highlights how hidden behavioral states may play a role in the dynamics of adaptive behavior. However, their model faces two essential limitations that are known to be unrealistic assumptions for biological systems, namely the discreteness of behavioral states and that transitions between states are memory-less (Markovicity) [26]. These two assumptions are helpful for avoiding the theoretical problems facing quantitative models of behavioral sequences. However, developing models that do not make those assumption can account for observed variation within behavioral states as well as account for observed context-effects [7].

## 3 Model Construction

As with the previous model, we take the units of our models to be the collective activity of neural circuits as depicted in Figure 2. We seek to develop systems of ODEs that describe the behavioral states. For pedagogical clarity and ease of visualization, we model the 3 behavioral states (forward, pause, reverse) rather than the 4 neural states (forward, inactive pause, reverse, active pause). We extend this model to the 4 neural states in the last section. We considered two different classes of non-linear dynamic networks, where each node in the network corresponds to one of the behavioral states in Figure 2. Using terminology from the literature on dynamical networks, we refer to each degree of freedom in our models as a *cell* [27]. The dynamics of each cell, *c*_*i*_, is defined by the following ODE:

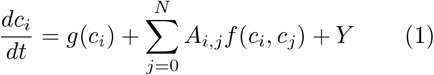

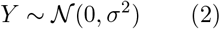

where *g* is the function representing the internal dynamics of a cell, *f* is the output function of a cell, and *A* is the adjacency matrix of the network. *Y* is a noise term which is used for some numerical simulations. *Y* is normally distributed around 0 with a standard deviation of 10^−1^ unless otherwise noted.

Networks of this form are commonly used to model complex systems in a wide range of domains [27]. An interesting property of such networks is that the homogeneity of the nodes implies the existence of equivariant subspaces [27]. Intuitively, we can observe that if cells are homogeneous, then swapping cells with the same connections should make no difference. The existence of equivariant subspaces leads to the existence of unique attractor dynamics that are not commonly observed in typical dynamical systems such as sequential state-switching [28], making them a promising class of model for understanding behavioral switching phenomena.

### 3.1 Sequence Model

We represent each element of the *C. elegans* behavioral sequence: forward, pause, and reverse, as the excitation of a cell in a dynamical network, setting *N* = 3 for equation 1. We are interested in choices of *g, f* and *A*, such that the resulting time series produces an activation pattern where the network of cells exhibits sequential dynamics. We consider dynamics to be sequential when the activity of the system shows small long-lived regions of state space and short-lived transitions between them. Since we take the activity of the cells to correspond to the behavioral modes, we also expect that these long-lived regions are such that the state of one cell is significantly higher than the state of the other two cells which are near the same low value. Examples of models that match this criteria are plotted in Figure 4.

**Fig. 3.**
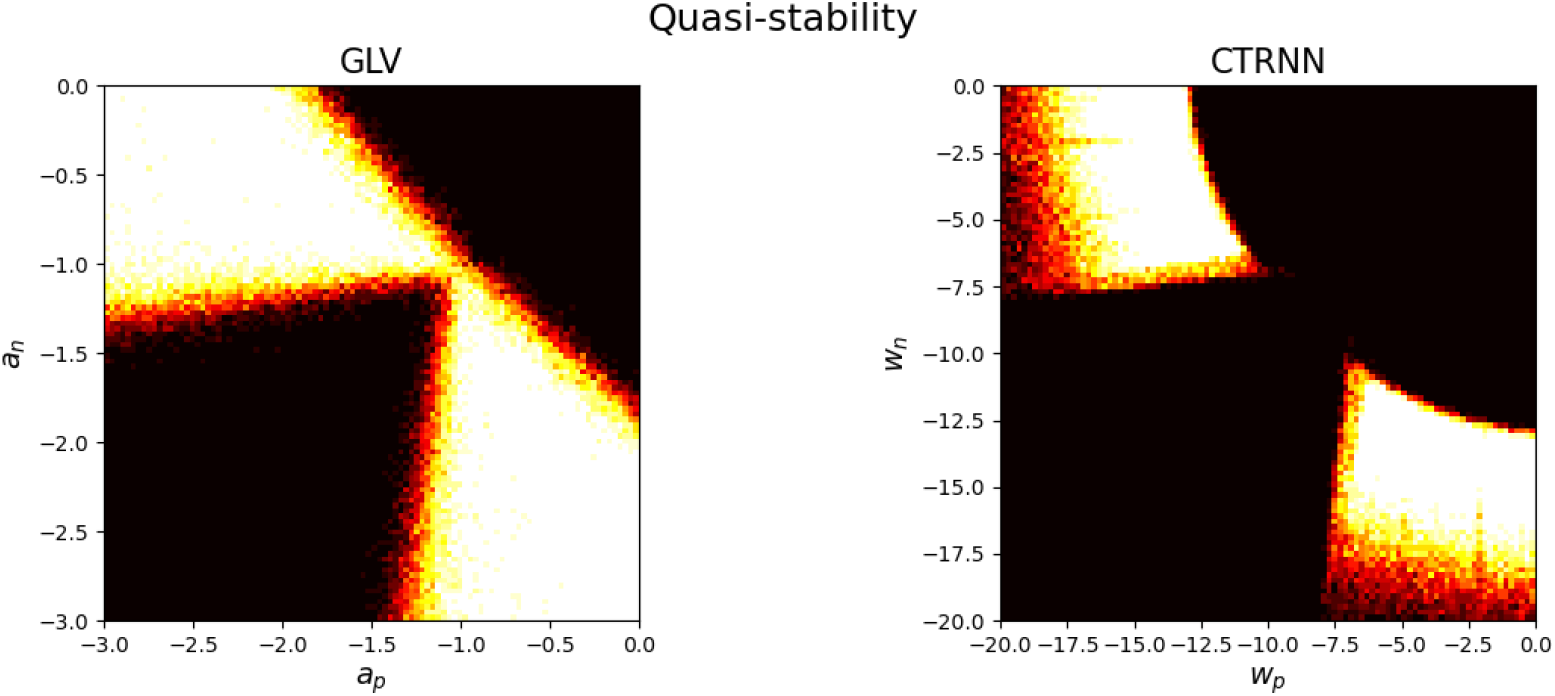
Heatmaps of parameter space for the two different dynamical models. Colored regions represent parameter combinations for which systems exhibit behavioral switching (defined in the text) regions in black are parameter combinations which do not result in behavioral switching.

**Fig. 4.**
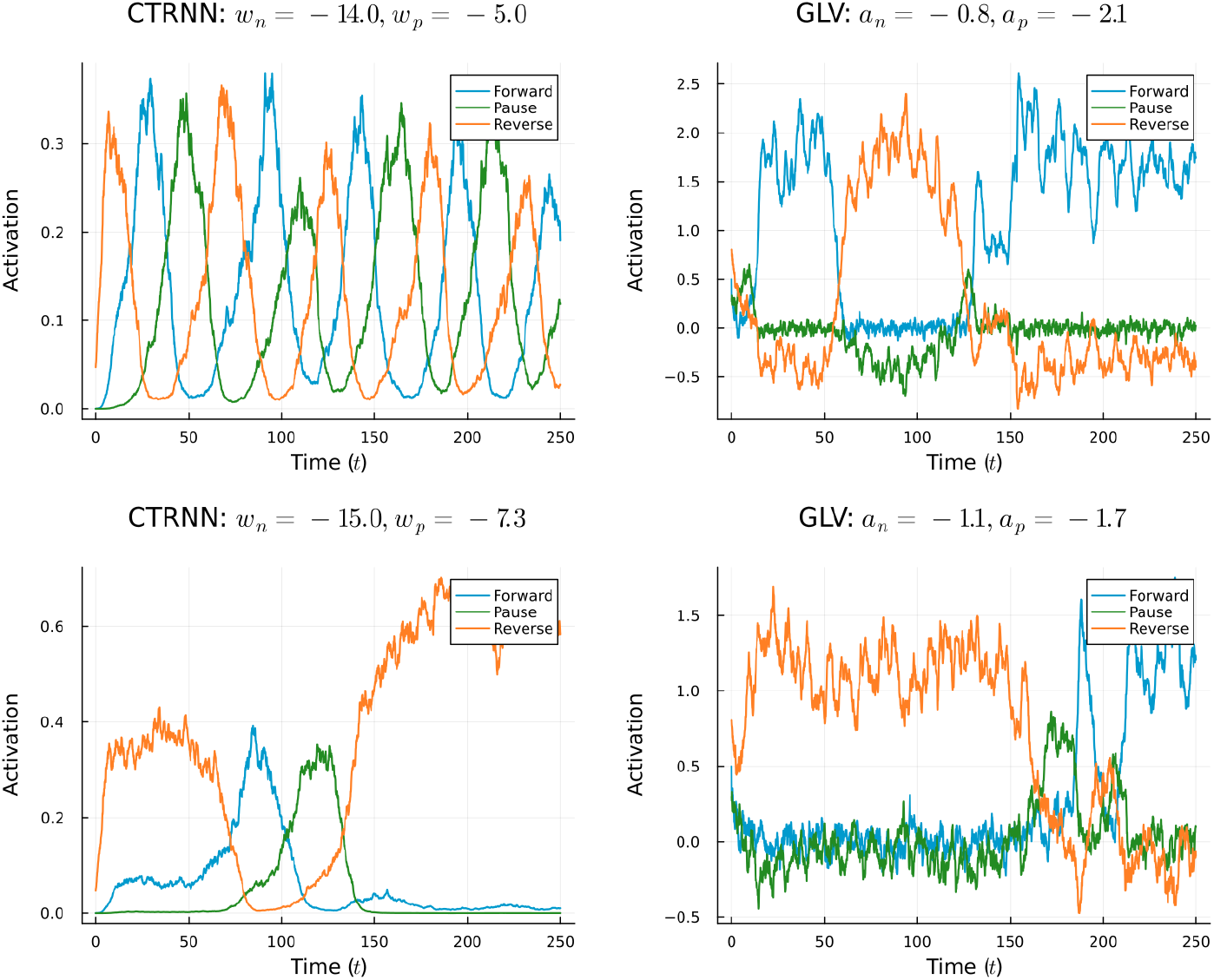
Example timeseries of parameters that exhibit behavioral switching.

The functions *g* and *f*, taken together, specify a class of dynamical networks for which different *A* matrices will give rise to a range of different dynamical landscapes. We used two different classes of dynamical networks: generalized Lotka-Volterra (GLV) models and continuous-time recurrent neural networks (CTRNNs). We chose these models because they have both been used to model the dynamics of cognitive states [29, 30] and because they are relatively well understood theoretically. We make use of this second condition in how we define the wiring matrix *A*.

Wiring dynamic network models to generate desired outputs is an open problem that does not have a generalized solution. However, various branches of nonlinear dynamics research have focused on how to relate functional dynamics to underlying network structure [27]. Significant progress has been made for various classes of dynamical networks, such as GLV [31] and CTRNNs [32]. Suited for our purposes, a branch of dynamical systems research focuses on construction methods for dynamical sequence generators [33]. These methods can be thought of as a set of rules that take as input a directed graph, in this case the graph in Figure 2, and return a set of conditions. If the adjacency matrix *A* obeys these conditions then it will exhibit sequential dynamics that embed the input graph. GLVs and CTRNNs are two such classes of dynamical networks for which such construction methods have been developed.

### 3.2 GLV Construction

The first construction method, proposed in [34], relies on the symmetries of an N-dimensional simplex where the corners of the simplex correspond to cells of the network, which also correspond to the dynamical states of the system. Dynamical transitions correspond to the convex edges between the points of the simplex. The simplex method was originally developed for the equation:

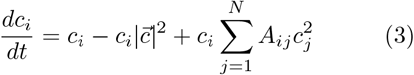

This system is governed by the linear combination of three terms. The first term is exponential growth of state *c*_*i*_. This growth is proportionally inhibited by the square sum of the activity of all cells, which can be interpreted as an energetic constraint that limits a cell’s growth to the interval (0, 1). The third component introduces the network connectivity structure with the adjacency matrix *A*.

The original simplex method states that given equation 3 and a desired graph, *G*, a matrix *A* can be defined such that an *N*-dimensional dynamical system can be constructed which contains a robust heteroclinic network that embeds *G*. The conditions for matrix *A* are as follows:

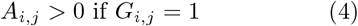

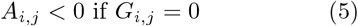

Intuitively, the first and second terms of equation 3 enforce a winner-take-all dynamic. In the case where *G* is unconnected, the matrix *A* simply adds further inhibition maintaining the winner-take-all dynamic. However, connections in *G* call for small excitatory connections from predecessor to successor states. This excitation breaks the global inhibition needed to maintain the dominance of the winning state. As a result the successor node will slowly increase its activity until it becomes able to compete with the dominating state, which causes a switch where the new node begins to dominate, a phenomena known as winnerlesscompetition [35]. We also note that this dynamic is only possible when the successor is guaranteed to overcome and inhibit its predecessor. This guarantee is only possible when there is no positive feedback between the successor and the predecessor node beyond this unidirectional excitation. This is not true in the case of cells that are their own successors or whose successors are also their predecessors. Hence, one- and two-cycles are not allowed in the input graph for this method. With the exception of one- and two-cycles, this method is quite expressive and allows for very complex networks of states to be embedded into relatively simple differential equations.

The previous equation can be rewritten into the form of Equation 1 by absorbing the global energy term into the adjacency matrix and taking:

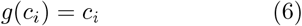

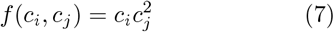

Although the original simplex method has a quadratic output function, it is known that the dynamics are closely related to the GLV system [36]. The only difference between these two systems is that:

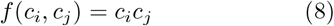

For the rest of this work, we consider this second variation because the GLV system has a long history in neuroscience [35, 37–39].

Putting equations 1 and 6 together, choosing *G* from Figure 2 and assuming symmetry between states, we can define a 3-dimensional system:

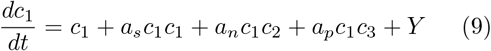

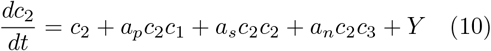

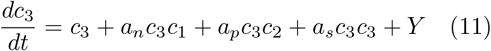

where *c*_*i*_ are the forward, pause, and reversal states. The proof of the construction method shows that we can set *a*_*s*_ = 1, leaving us with only two degrees of freedom: *a*_*p*_ and *a*_*n*_ (the connection from the previous cell and the connection from the next cell respectively).

### 3.3 CTRNN Construction

The second class of models we consider is the CTRNN (equivalently, Wilson-Cowan [40]) neural mass model. CTRNNs are commonly used as firing-rate models that have been applied in a wide range of neural contexts [41] and have been studied in some theoretical depth [42]. CTRNNs have proven a useful playground for better understanding concepts that appear in more complex neural models [43] while remaining relatively tractable using standard numerical techniques. In [40], the authors showed how networks of fixed points could be constructed in CTRNN dynamics, making them a useful candidate model for understanding metastability and behavioral switching. CTRNNs are defined by taking:

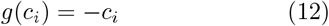

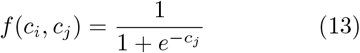

In this case, each cell possesses an inherent decay term rather than a growth term, and the output function is a monotonic, bounded sigmoid function rather than a quadratic. As a result, unlike the GLV-type equations, CTRNNs do not possess the same strict symmetries. This changes the kinds of metastability they can exhibit. Namely, structurally stable heteroclinic dynamics cannot occur in CTRNNs because they lack an equivariant symmetry. We will further discuss the implications of this property in the section on bifurcations.

The conditions for the CTRNN construction method cannot be written as succinctly as the simplex method, and we refer the reader to the appendix, in section 9, for the necessary conditions. We also note that this method suffers from similar restrictions as the simplex method: it does not allow for self-loops or two-cycles in network connectivity. Additionally, due to intrinsic properties of CTRNNs, this method also does not allow for *δ*-cliques, a pattern in which there is a skip connection between two states that are removed from each other by one step in the sequence.

Making use of the graph of our system in Figure 2, we end up with the equations:

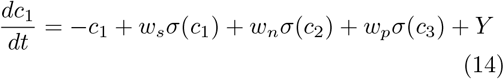

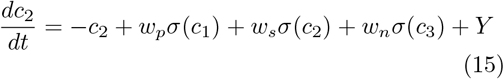

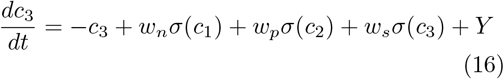

where *Y* is the same as in the GLV case, using the construction method for CTRNNs, we can set *w*_*s*_ = 1, again leaving us with two degrees of freedom for parameterization: *w*_*p*_ and *w*_*n*_. Where *w*_*p*_ is the input from a predecessor state to the successor state and *w*_*n*_ is the input from the successor state to the predecessor state.

### 3.4 Method Comparison

Applying the above construction methods resulted in two different systems of equations, each with two degrees of freedom for their parameters and three phase dimensions. We sought to find parameters such that the resulting dynamics generate the behavioral states laid out in Figure 2. To determine whether a given model generated the 3 desired behavioral states and transitions quickly between them, we performed a scan across the two parameters in both systems.

To calculate which networks match these criteria, we simulated models for 500 time units. We then cut off the first 100 time units as an initial transient. For each time unit, we determined whether there was a dominant state by checking if any of the three cells had an activity level greater than the sum of the other two. We then calculated the length of time that the state was dominant. We considered a system to exhibit quasi-stability when the length of dominance time was at lease twice the length of transition time. We assigned a score to each system proportional to the number of dominant states it exhibited that match these criteria. We required that a system must exhibit all 3 states as dominant and must exhibit at least 4 transitions between states. We plotted the results averaged over 10 runs per parameter pair in Figure 3.

All models were numerically integrated using the Julia DifferentialEquations package [44], with simulations performed using the Euler-Maruyama method. We generated contour plots, depicted in Figure 3. To determine the valid parameter ranges for which we observe the desired behavioral sequencing behavior. We also plotted two example time series for each model class as representatives for the dynamics of individual models in Figure 4. All plots are plotted in the output space of *f* (*c*_*i*_), therefore the artifacts of symmetries in GLV case are ignored and the resulting dynamics are bounded between (0,1) in the CTRNN case.

From the resulting time series, we saw that both models were capable of producing activation sequences that match the observed empirical data. The major difference between the two models is the state of the inactive cells. In the case of the GLV equations, the inactive cells have a value near 0. While in the CTRNN equations, the predecessor cells show slight activation. Given that we were able to successfully model the target system, we sought to explain the mechanisms underlying the observed model behavior.

## 4 Dynamical Mechanisms of Behavioral Switching

We found that for a range of parameters, both classes of models were capable of reproducing the observed sequential phenomena. We took a deeper look at their dynamics to better understand these two model classes and their differences. In this section, we explicate the differences between the models by considering their underlying phase space structure. This will also give us further insight into the more general features of behavioral switching.

We started by identifying the ranges of parameters for which we observe behavioral switching. These are depicted in Figure 3. These heatmaps are symmetric due to the parameter symmetries in *a*_*p*_(*w*_*p*_) and *a*_*n*_(*w*_*n*_). The difference between them is the order of the state switching. When noise is present, switching can deviate from strict ordering [45], but, in the deterministic case, we observed distinct orderings. For this section, without loss of generality, we pick the set of parameterizations where *a*_*p*_ *< a*_*n*_ and *w*_*p*_ *< w*_*n*_ which corresponds to an ordering where the pause state occurs after the forward and before the reverse state.

From the deterministic dynamics, we identified that the models were relying on three distinct possible dynamical mechanisms for behavioral switching:

- Attractor Hopping: characterized by fixed-point deterministic dynamics.
- Heteroclinic Channels: characterized by an oscillation with a growing period.
- Haunted Limit Cycle: characterized by an oscillation with a constant period.

For each of these classes of solutions, we were interested in what dynamical structures give rise to the occurrence of the individual quasi-stable behavioral states and what dynamical mechanisms were responsible for the transitions between them. We used the Dynamica package in Mathematica for our analysis.

### 4.1 Attractor Hopping

The first class of solutions we identified occurred both in the CTRNN and GLV model equations. This class of solutions eventually exhibit equilibration for the majority of their initial conditions when noise is removed. The resulting equilibrium corresponds to one of the three behavioral states. The behavioral states to which the system converges depends on the initial conditions. This shows that the system is multistable. To investigate properties of these systems, we performed a phase space analysis (Figure 6).

**Fig. 5.**
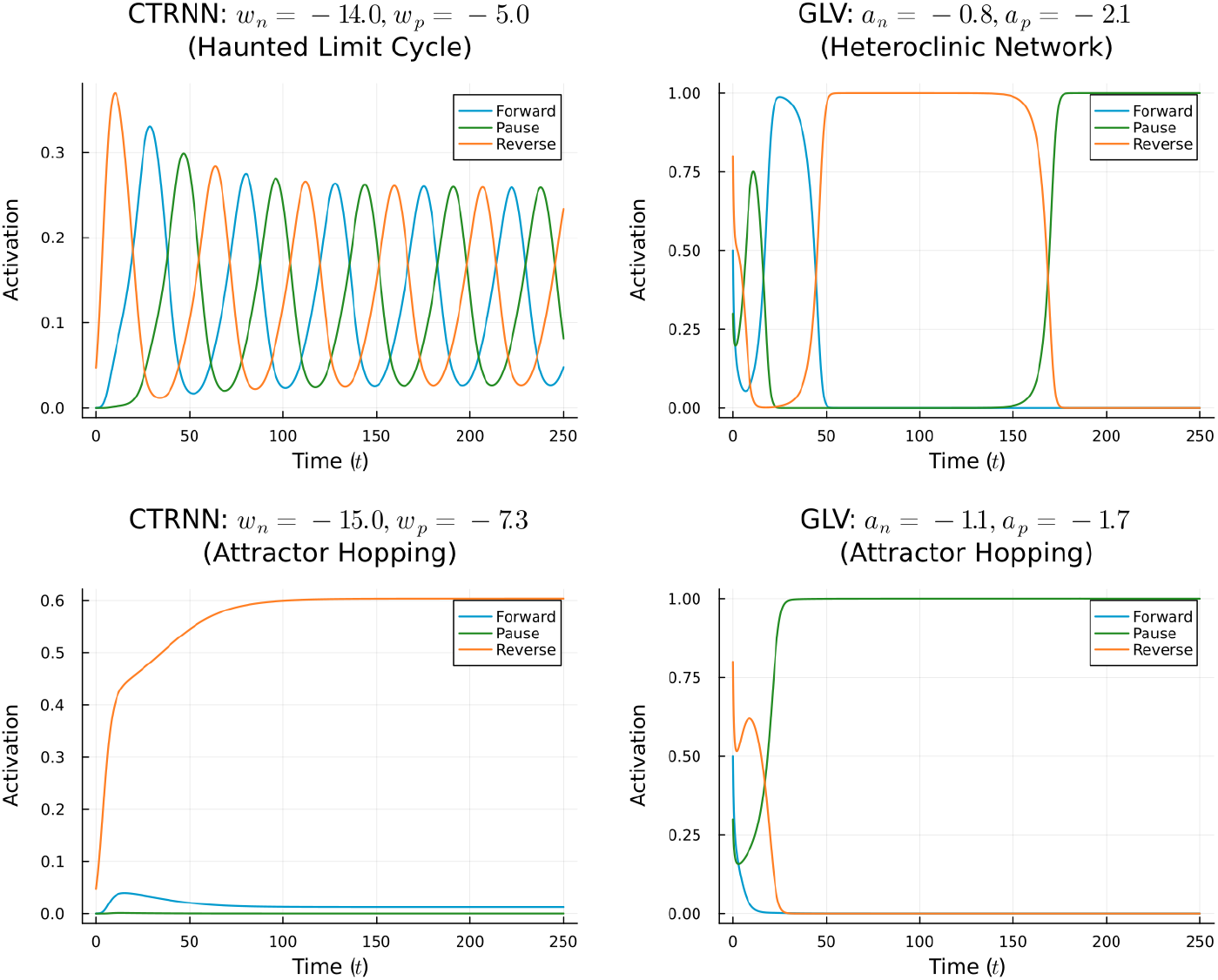
Deterministic components of the same models that generated the time series in Figure 4.

**Fig. 6.**
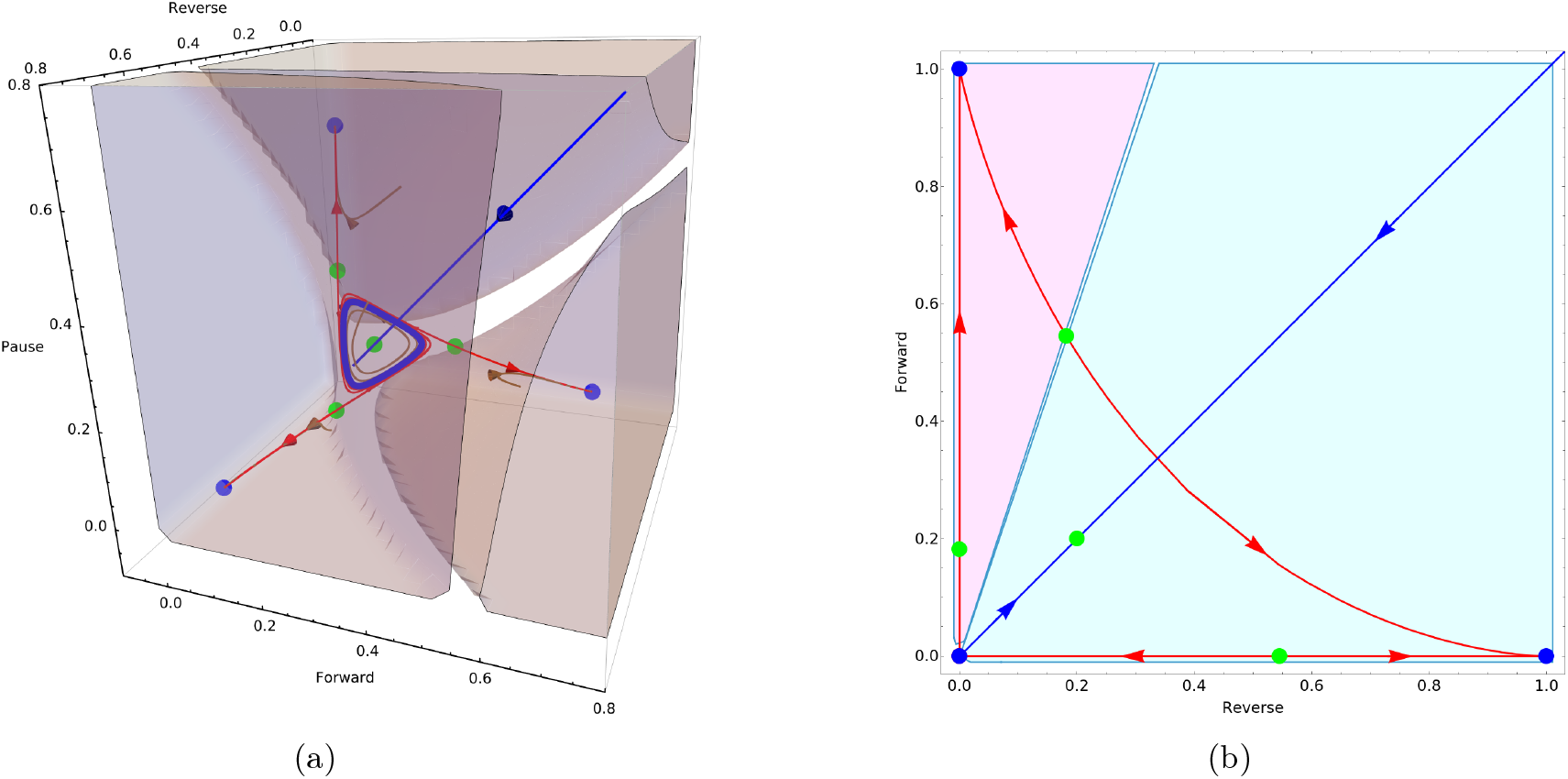
(a) Phase portrait of the CTRNN system that produced that timeseries in Figure 5. Attractors are plotted in blue including the individual point attractors that correspond to behavioral states as well as the limit cycle near the quiescent state that is not observed in the timeseries. Saddle points are plotted in green. The brown regions represent the basin boundaries of the individual behavioral states. (b) Two dimensional projection of the GLV system that produced the time series in Figure 5. This projection visually demonstrates how the saddle state divides the boundaries of the basins of attraction. The forward and pause basins are plotted here in magenta and cyan respectively. All other colors correspond to the same convention as in (a).

We first applied fixed point analysis to the CTRNN model by finding solutions where 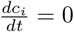 for all *i*. This process resulted in the identification of several fixed points, though not all were attractors and thus not candidates for behavioral states. To identify the attractors, we performed stability analysis. Since we restrict ourselves to point attractors, this analysis is straightforward: we linearize the system at each fixed point and examine the eigenvalues. Stable fixed points have all negative eigenvalues, unstable points have all positive eigenvalues, and saddle points have mixed eigenvalues.

Plotting these fixed points colored by stability in Figure 6, we observe three stable fixed points corresponding to behavioral state saturation. The stable fixed points are separated by saddle points for which the stable manifolds act as a separatrix. At the center of phase space, we find an additional fixed point representing the quiescent state of all three neural masses. We found that in most solutions, the quiescent fixed point is a saddle. Plotting trajectories near the saddle quiescent state, we observed oscillatory behavior, suggesting that a small amplitude limit cycle was present. We used a shooting method to explicitly identify the limit cycle and plotted it along with the equilibria in Figure 6. In a small subset of solutions, we found that the quiescent state was a stable fixed point without a surrounding limit cycle.

We then applied the same analysis to the GLV models that exhibited similar equilibrium dynamics. The resulting phase diagram showed that the stable fixed points similarly corresponded to the active states of the individual cells. Between each pair of fixed points, we again found saddle points each with its stable manifold acting as a separatrix dividing the basins of attraction. The solution at the center of the cycle was also a saddle point. At the origin we found a non-hyperbolic fixed point serving as a singularity from which new invariant sets may emerge. The key differences from the CTRNN system are: the unstable nature of the origin in GLV (versus the stable quiescent state or stable small amplitude limit cycle in the CTRNN), and the presence of a non-hyperbolic singularity in GLV that could generate new dynamical invariants.

Even though we identified three different types of phase portraits (with and without limit cycles in the CTRRNs and with a singularity in the GLVs), we consider all these models to exhibit the same dynamical mechanism for behavioral switching. The basic mechanism for this class of solutions is that the system settles into one of the attractors corresponding to the peak activation of one cell; perturbations due to noise cause the system to wander around the basin of attraction of that behavioral state; over time, the system meanders across an attractor basin boundary which leads to behavioral switching.

However, not all multistable parameterizations of the CTRNN and GLV system result in switching dynamics. For switching to occur, the stable manifold of the separatrix must be close enough to the equilibrium so that the noise can tip the system into another basin of attraction by crossing the separatrix. Both of the models exhibit parameter regimes for which this is indeed the case. The role of the saddle manifold and how it divides the basin of attraction can be seen in the Figure 6 where we plot the two dimensional projection of the GLV dynamics showing how the stable manifold of the saddle point divides the phase space.

### 4.2 Heteroclinc Connections

The majority of the solutions that we found in the GLV system did not make use of the attractor hopping mechanism. Most of the solutions, when noise was removed, exhibited an oscillatory-like behavior. This behavior is not exactly oscillatory because the oscillations show a growing period with fixed amplitudes. The saturation of individual cells lasts longer and longer on each iteration. This is a known signature of a special class of dynamics called heteroclinic dynamics.

Similar to our previous analysis, we solved the system numerically to find equilibria and linearized around them to determine stability. We found a number of fixed-point solutions, none of which were attractors, despite the system exhibiting clear quasi-stability in its dynamics (Figure 7).

**Fig. 7.**
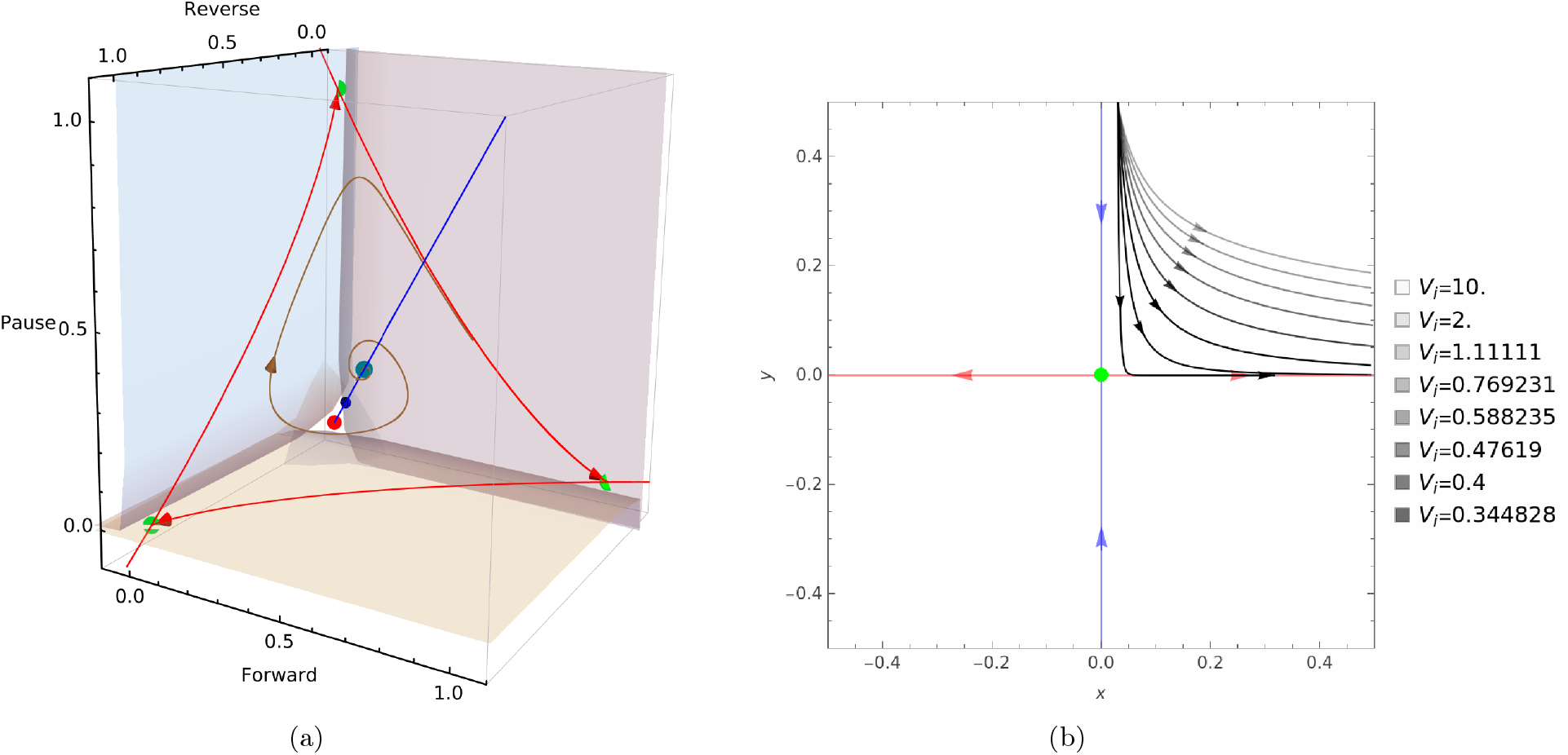
(a) Phase portrait of the GLV system that generated the time series in Figure 5. Saddle points are colored green and tinted blue based on their stability and the unstable equilibrium is colored in red. Unstable manifolds of the saddle points are plotted in red and stable manifolds are plotted in blue. The planes are the stable manifolds of the saddle points that correspond to the behavioral states. An example trajectory is plotted in brown. (b) A simple saddle point with an unstable manifold in red and a stable manifold in blue. Individual trajectories plotted in black correspond to different saddle values with lower opacity corresponding to smaller saddle values.

Instead, we identified saddle and unstable fixed points in the phase portrait. We could easily ruleout the unstable equilibrium point as a potential cause of the behavioral states. This left the saddle points as the source of the quasi-stable states. However, we observed 4 which implied not every saddle correspond to a behavioral state. To determine how the saddle points were contributing to the occurrence of the behavioral states, we had to make use of an additional concept from the toolkit of dynamical systems theory: the saddle value.

Given a saddle point one can find its saddle value based on the ratio of it its (absolute) smallest negative eigenvalue, *λ*_*min*_ and its largest positive eigenvalue *λ*_*max*_. Explicitly:

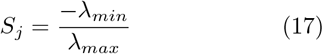

Where *S*_*j*_ is the saddle value for the saddle point *j*. We plot the dynamics of a single 2-D saddle point in 7b to provide intuition. Put simply, the strength of the saddle value determines how strongly a trajectory will approach the stable manifold of a saddle point before being pushed onto its unstable manifold.

Metastable states are characterized by saddle values greater than 1, indicating dissipative saddles which trajectories approach over time. The saddle value can also be used to calculate the stability of a heteroclinic network as a whole by taking the product of all saddle values in the network; a heteroclinic network is stable if and only if this product is less than 1 [46]. Coloring nodes by stability (and saddle value for saddles), we observe the state space structure in (Figure 7a). Four dissipative saddles are visible, with one acting as an entry point, compressing trajectories onto the metastable subspace, this is a feature common in heteroclinic dynamics [47].

Heteroclinic networks enable state transitions without external perturbations, distinguishing them from excitable networks. Their defining feature is the alignment of stable and unstable manifolds between nodes: the unstable manifold of one node aligns with the stable manifold of another, creating cyclical dynamics. In the case of our GLV system in Figure 7a, the unstable manifold of a cell, in red, lies in the stable manifold of the next cell, the blue planes. This leads to a phenomenon known as *winnerless competition* [48] which arises from local excitation and long-range inhibition, where neural regions temporarily dominate one another. In deterministic contexts, the characteristic of a heteroclinic time series is a fixed amplitude oscillation with increasing period as in Figure 5. However, when noise is present trajectories meander around the saddle points rather than approaching them directly, this leads to development of an average period and hence makes the system resemble more traditional cyclic behavior.

### 4.3 Haunted Limit Cycle

Unlike the attractor hopping and heteroclinic channels, the metastable dynamics in the CTRNN system cannot be explained through standard invariant analysis. When behavioral states correspond to explicit invariants (e.g., fixed points or limit cycles), numerical solving and continuation methods can directly identify and characterize them. However, for the CTRNN’s metastable regime, this approach fails.

Performing standard fixed-point analysis reveals a single saddle equilibria, which does not align with the system’s observed slow states (Figure 5). The deterministic trajectories show a clear oscillation and when plotted in phase space, these oscillation surround the saddle point. This is suggestive that these trajectories are approaching a non-equilibrium invariant solution: a limit cycle. Using a shooting method we were able to identify the exact position of this limit cycle plotted in Figure 8a in blue.

**Fig. 8.**
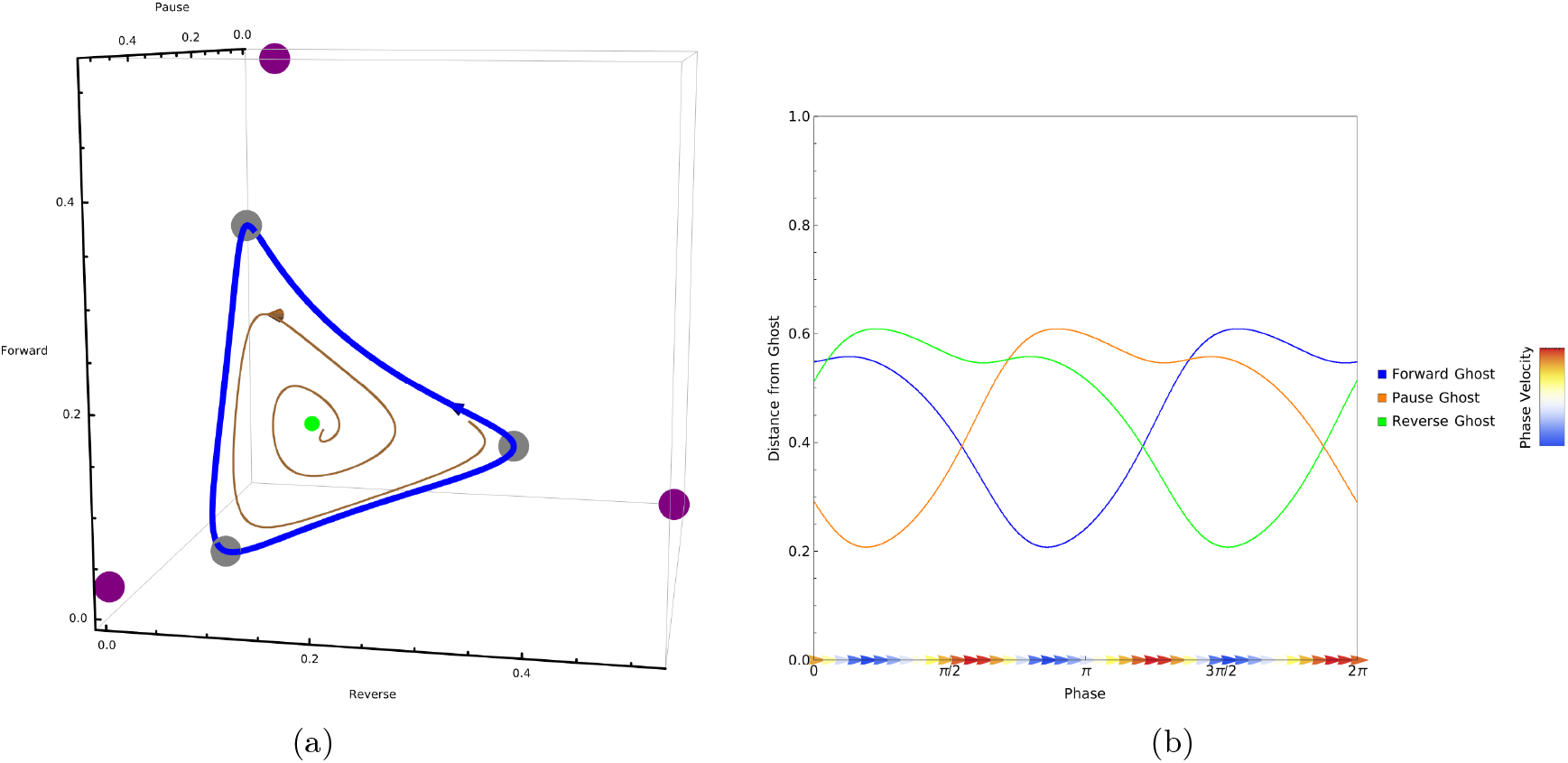
(a) Phase portrait of CTRNN system that generated the time series in Figure 5. Stable limit cycle is depicted in blue. The quiescent saddle point is depicted in green. An example trajectory is plotted in black. Gray spheres correspond to slow points along the limit cycle. Purple spheres correspond to the location of the real components of the conjugate solutions in the complex plane. (b) Magnitude of the velocity along the limit cycle. Points plotted at the slow points along the limit cycle with the y-component representing the distance of the point from the ghost in the complex plane.

These two invariant solutions fully characterize the long-term behavior of the system across all initial conditions. However, we were interested in not only characterizing long-term behavior but explaining why this system exhibits the occurrence of quasi-stable behavioral states which manifested as slow points along the limit cycle. To find the exact location of these slow points, we considered the velocity of the system trajectories along the phases of the limit cycle (Figure 8b). These slow points correspond to the behavioral states of our system. The mechanism causing these slow regions along the limit cycle could not be explained using standard phase portrait analysis.

We hypothesized that slow-point behavior emerged from a different kind of dynamical phenomenon known as ghost states [13]. Ghost states are slow regions inside the vector field which are caused by the disappearance of equilibria after a bifurcation. In ODEs, when a fixed point disappears the derivative at the point which was previously 0 at that point is still a local minima (as a result of continuity). Ghost states are absent from the real vector field but can be inferred through the dynamics. Recent advances now allow explicit design of ODEs with ghost states of desired topology [40], but their analysis requires advanced techniques [49].

A simple method for identifying ghost states is to explicitly calculate the velocity of the dynamics in phase space and identify local minima where the system exhibits slow dynamics. An alternative means of identifying ghost states, in simple cases, is via analytic continuation into the complex plane [13]. One can extend fixed-point identification by using complex initial conditions and analytically continuing the equations to operate in complex space. This approach works for point-like ghost states but becomes intractable for geometrically complex invariants (e.g., ghost limit cycles or tori). Stability analysis of such structures remains an open problem in criticality research.

Plotting both real fixed points and identified ghost states (Figure 8), we see that while metastable states don’t coincide with equilibria, the latter still shape the dynamics. In particular, although the oscillatory behavior we observe in the dynamics is due to a limit cycle, this limit cycle is haunted by the ghosts of a bifurcation to come that will lead the system to have equilibria at the slow points.

Quantifying haunted invariants remains challenging. Both ghost-state identification and the catalog of invariants they can haunt are open questions. Unlike *winnerless competition*, ghost-state transitions need not involve dominance by another state; a ghost may vanish independently, leaving no trace in the real-plane dynamics.

## 5 Bifurcation Analysis

Having identified the different mechanisms underlying the various models, it seemed that we were observing very different structures not only between different model classes but even within model classes. To get a better sense of how the same architecture gave rise to such different behaviors, we turned to bifurcation analysis. Here again, we used the Dynamica Mathematica package [50].

It is known that heteroclinic connections and haunted-invariants differ topologically [15]. Heteroclinic cycles rely on saddle connections, while ghost cycles emerge from a destabilized attractor. However, bifurcation analysis reveals their shared origin in the collapse of multistability, exposing a deeper continuity. Both regimes arise via codimension-one bifurcations that destabilize attractors, redirecting trajectories along metastable pathways. Both the GLV and CTRNN systems exhibit a transition from multistability to metastability guided by the interaction of a Hopf bifurcation and a fold bifurcation. This shared bifurcation structure coincides with a conserved ordering. The heteroclinic phase lies between multistable and ghost phases, regardless of the measure of the parameter regions they occupy.

### 5.1 Bifurcations fo the GLV System

Bifurcations of the 3D GLV system have been analyzed previously in [51]. Based on these previous results, we focus on parameter ranges relevant to the network dynamics, *a*_*p*_, *a*_*n*_ ∈ (0, 3). Our results revealed three key bifurcation curves:

1. A diagonal Hopf bifurcation curve along *a*_*p*_ = −*a*_*n*_ + 2, identified by tracking the central attractor. We also note that this is a degenerate Hopf bifurcation that is atypical due to both its co-occurence with a fold bifurcation and due to the enforced parameter symmetries.
2. Two fold bifurcation curves at *a*_*p*_ = 1 and *a*_*n*_ = 1, we note that in fact these two bifurcation curves are actually each three overlapping bifurcation curves thus the system displays 6 fold bifurcation curves in total. This overlap is the result of our symmetry assumption since the fold bifurcation occurs in each behavioral state at the same parameter values.

The bifurcation curves intersect at *a*_*p*_ = *a*_*n*_ = 1; however, the intersection does not constitute a higher co-dimension bifurcation because these bifurcations occur at different points in phase space. These bifurcations partition the (*a*_*p*_, *a*_*n*_) parameter space into three distinct stability regimes (Figure 9):

1. **Central Fixed Point Dominance**: For *a*_*p*_ *<* 1 and *a*_*n*_ *<* 1, the system exhibits a single stable fixed point at the coexistence equilibrium. This regime corresponds to the region bounded by the two fold curves, where mutual inhibition becomes supercritical.
2. **Multistability**: Crossing either fold bifurcation curve (*a*_*p*_ *>* 1 and *a*_*n*_ *>* 1) Beer2021Dynamicaintroduces bistability between dominant states. Here, the predecessor and successor populations act as mutual “switches,” with hysteresis governed by the fold bifurcations.
3. **Heteroclinic Dynamics**: Near the intersection point at (*a*_*p*_, *a*_*n*_) = (1, 1), the interplay of Hopf and fold bifurcations creates two regions supporting heteroclinic orbits. These manifest as slow switching dynamics between states when parameters approach the Hopf line from either side.

**Fig. 9.**
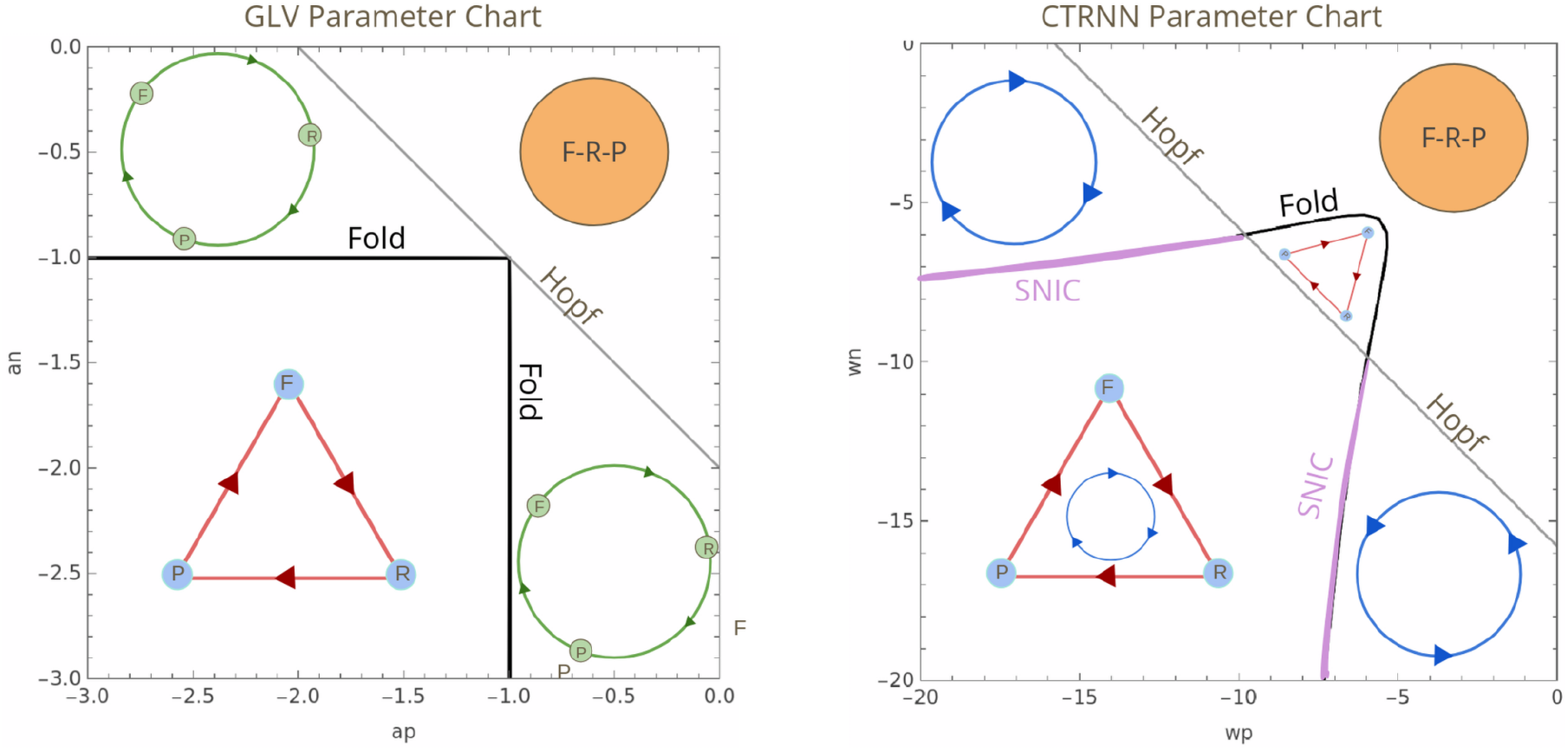
Parameter charts for the GLV and CTRNN systems. The region with a single stable fixed point is denoted with a filled orange circle where the forward, reversal, and pause cells are equally active. Regions that exhibit heteroclinic network are denoted by a green circle. Regions exhibit haunted limit cycles are denoted by blue circles. Regions with multistable dynamics are denoted by a red triangle.

We note that the degeneracy of the Hopf bifurcation has interesting implications. In particular, exactly along the bifurcation curve the system exhibits a non-hyperbolic limit cycle. Properties of such structures, including within the context of GLV equations, is a topic of mathematical interest [52].

The intersection point acts between the bifurcation curves as an organizing center, mediating transitions between these regimes. Metastability emerges along the Hopf curve, where damped oscillations about the central fixed point coexist with slow transitions between saddle states. This structure aligns with earlier findings while clarifying how bifurcations interact to give rise to a heteroclinic structure.

### 5.2 Bifurcations of the CTRNN System

The bifurcation space of CTRNNs has been previously explored in [32, 42]. Here we are interested not in the general bifurcation structure, but in the bifurcation structure induced by the same symmetry applied to the GLV system as in equation (5). Our results revealed two key bifurcation curves:

1. A diagonal Hopf bifurcation curve along the *w*_*p*_ = −*w*_*n*_ + 2 line, identified by tracking the central attractor.
2. A fold bifurcation curve, located by following attractors at their corresponding states. This fold bifurcation within the relevant parameter range (above the *w*_*p*_ + *w*_*n*_ = 1 line) also occurs along an invariant cycle in phase space. Therefore in this region the bifurcation is an instance of a special class of fold bifurcations known as saddle-node on an invariant cycle (SNIC) bifurcations which can create or destroy limit cycles.

These bifurcations partition the (*w*_*p*_, *w*_*n*_) parameter space into four distinct stability regimes (Figure 9):

1. **Central Fixed Point Dominance**: In the lower left region, the system exhibits a single stable fixed point at the coexistence equilibrium. This regime corresponds to the region bounded below both the fold curve and the Hopf bifurcation, where mutual inhibition remains supercritical.
2. **Multistability w/o limit cycle**: Crossing the fold bifurcation curve introduces bistability between dominance states. Here, the predecessor and successor populations act as mutual “switches,” with hysteresis governed by the fold bifurcations, similar to the GLV system.
3. **Multistability w/ limit cycle**: Across the (SNIC) bifurcation curve and passed the Hopf bifurcation is a region similar to the other multistable region except that in this region the separatrices occur along the invariant cycle. This also means that the system expresses small amplitude oscillations for a small subset of initial conditions.
4. **Haunted Limit Cycle**: In the subcritical region of the Hopf curve but before entering the supercritical region of the fold bifurcations, the interplay of the Hopf bifurcation and the approaching fold bifurcations creates two regions supporting limit cycles which are haunted by the ghosts of the attractors. These manifest as slow switching dynamics between states when parameters approach the Hopf line from either side.

The system is very similar to the GLV system in that its bifurcation structure is governed by the point at the center of the cycle, giving rise to oscillatory dynamics, and the fold bifurcation occurring at the excited state of each neuron. The competition between this cyclic behavior and the dominance of the individual nodes gives rise to metastable dynamics that resemble, but are not identical to heteroclinic dynamics.

We also note that along SNIC/Fold bifurcations, the CTRNN system exhibits a set of nonhyperbolic saddle points. This implies that at the exact point at which the SNIC bifurcation occurs, the system exhibits a non-hyperbolic heteroclinic cycle. This structure is known to be related to the occurrence of ghost cycles [53]. This is also an interesting counterpart to the non-hyperbolic limit cycle exhibited by the GLV system.

## 6 Matching Dwell Time Distributions

So far, we have only considered the sequential ordering of the behavioral states as outlined in Figure 2. However, behavioral states are not all equivalent. Some behaviors take longer than others and it has been shown that in the case of *C. elegans* foraging, this difference is in fact functional. The time dwelt in the behavioral states determines the strategy being implemented [23] by a foraging worm. To account for this, we sought to incorporate heterogeneous dwell times into our models. To add to biological relevance, we used the four-state neural model instead of three behavioral states. We use data from [8] which includes the mean dwell-times of the states in Figure 2. The empirical dwell-times are plotted in Figure 11.

To construct the four node models we again used equations (1) and (2), this time with *N* = 4. We used the adjacency matrix based on the neural states in Figure 2. The construction methods for both the GLV and CTRNN systems could still be applied. However, the resulting sets of differential equations do not determine the actual dwell time for the behavioral states.

Adjusting the dynamical equations in (5) and (3) to accommodate for heterogeneous dwelltime distributions is non-trivial and requires an understanding of the stability properties of the equations. Since both the GLV and CTRNN equations are nonlinear, small changes to the weights can have a significant impact on the resulting dynamics and although our bifurcation is informative about the possible kinds of behavior it is not sufficient to determine the appropriate edge weights for the desired dwell-times. To appropriately parameterize the model we had to turn to a different set of techniques

### 6.1 GLV Dwell Time Control

Controlling GLV dwell time dynamics has been addressed previously in [46] for the case of cyclic GLV systems and more recently for GLV systems with splitting transitions [54]. The primary insight for this approach is that for a GLV system under a given noise-intensity, one can calculate the mean dwell-time at a given equilibrium. Further, this quantity can be approximated given that the parameters of the system are known.

In [46], the authors demonstrate that the dwell time of a state in a noisy heteroclinic network can be cast as a first-passage time problem. Along with some simplifying assumptions and approximations, they show that the mean dwell-time can be written as a function of the GLV parameters. Rewriting this function by taking the parameters to be the unknowns and the desired dwell-time to be specified, a formula for the parameters can be written as:

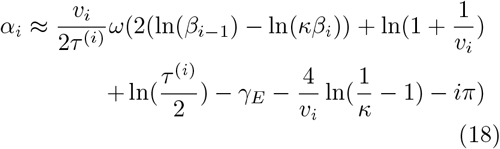

Where *τ* is the desired dwell time, *γ*_*E*_ is the EulerMaschoroni constant, *v*_*i*_ is the stable eigenvalue at node *i*, and 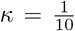 is chosen empirically. We also note that in our terminology from equation (1) this approach requires us to modify *g*(*c*_*i*_) by adding a constant multiplier *α*. This *α* is needed to counterbalance the change in stability caused by the modification of the weights between cells. For a detailed explanation of this equation and its derivation, we refer the reader to Appendix 1.

Solving the equation above for the desired dwell times we get the GLV equations in (**??**).

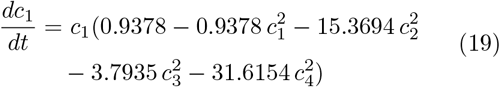

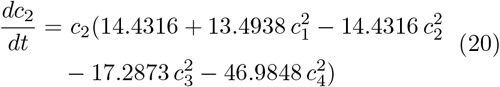

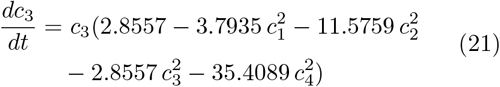

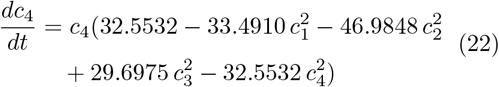

### 6.2 CTRNN Dwell Time Control

While we could explicitly approximate the appropriate parameters for the GLV system, no such approximation for CTRNNs has yet been developed. As a result to appropriately parameterize the CTRNN model, we performed stochastic search using an evolutionary algorithm [55]. To use stochastic search we had to first define an objective function that would result in the desired dyanmics. We defined the objective function to give a positive value if three conditions were met

1. The system exhibited individual behavioral states, meaning that we rewarded solutions which exhibit phases where a single cell’s activity is greater than the activity of other two cells. We weighted the reward by the amount of activity difference.
2. The dwell times within a given state correspond to the experimentally observed dwell times in [8]. Where we measured average dwell time and took the magnitude of the difference between the average dwell time and desired dwell time. We assigned high scores to solutions that minimized this value.
3. The system preserved the sequence as defined by the model in Figure 2. We rewarded solutions that preserved the sequencing defined in the Roberts et al. model.

We seeded a population of 1000 and evolved them over the course of 1000 rounds. The genome of the agents specified the weights between cells as a well as an additional gain term. Similar to the GLV case, we needed to add a parameter in the equations to account for the change in stability. We used a gain term, *g*_*j*_, which is similar to the growth rate *α* in the GLV model, it modifies the function *f* (*c*_*i*_, *c*_*j*_) = *σ*(*c*_*j*_ ∗ *g*_*j*_). Writing out the equations for the best agent after the evolutionary run, we get:

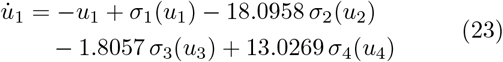

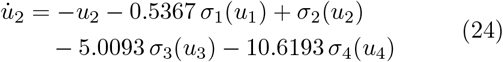

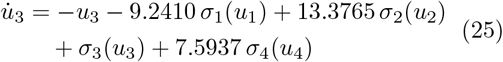

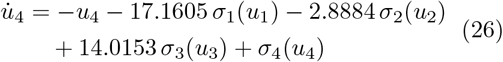

where each sigmoidal activation is:

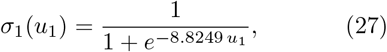

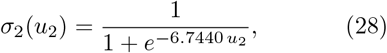

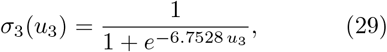

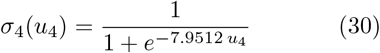

### 6.3 Model Performance

To assess the model performances, we simulated the resulting models using numerical integration and a noise term with variance 10^−1^. We then simulated the networks for a period of 500 time units across 10 runs. For each run we determined the dwell time for each of the 4 behavioral states. We then plotted the mean and variance of the models along with the empirically observed dwell-times in [8] (Figure 10).

**Fig. 10.**
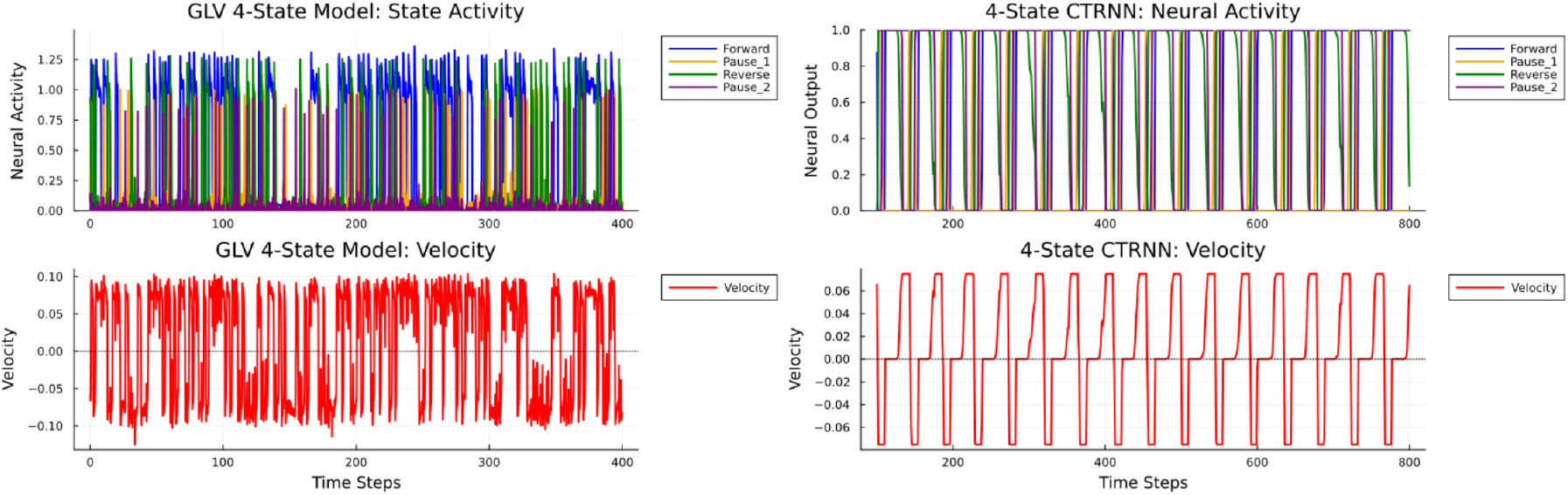
Two examples of simulated stochastic dynamics for GLV system (left) and CTRNN system (right). Velocity is calculated by subtracting the excitation of the reversal cluster from the excitation of the forward cluster and multiplying the result by the maximal forward speed of the worm based on the data from [16].

**Fig. 11.**
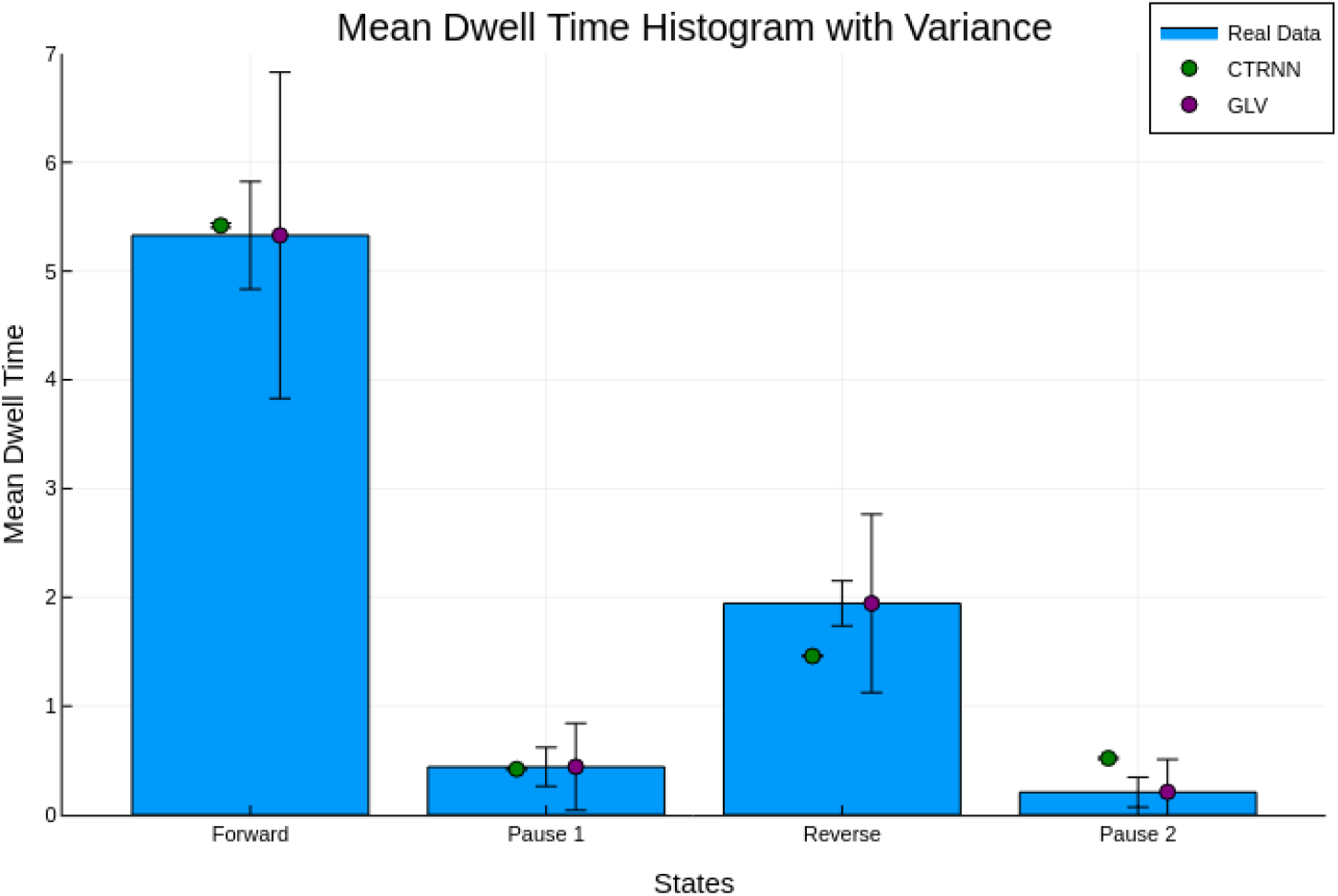
Comparison of models to real data from *C. elegans* behavior. Each point represents the mean dwell time for each model based on an integration over 1600 time units.

The results demonstrate that both models can closely match the measured observables. However, we notice that the spread of measured dwell times is significantly smaller for the CTRNN network than the GLV network even at larger noise levels. This is another feature that distinguishes haunted limit cycles from heteroclinic networks [49].

## 7 Discussion

In this work, we developed several different models of the forward-reversal in *C. elegans*. We first showed how two classes of equations, Generalized-Lotka Volterra (GLV) and continuous-time recurrent neural networks (CTRNNs) could be used to model sequential switching dynamics similar in application to Markov chains, however, without assuming Markovian properties between behaviors. We then used simplifying symmetries to expand on the different dynamical structures that play a role in the behavioral switching. We also used bifurcation theory to demonstrate how these dynamical structures relate to one another, revealing a shared underlying symmetry-breaking. Finally, we demonstrated how more complex models can be constructed by optimizing parameters to match empirical data.

In this work, we limited our consideration to models of quasi-stability where the quasi-stable states are conceived of as points. This is an unrealistic assumption since behavioral states are not points in states space but rather complex dynamical structures. However, this choice allowed us to significantly reduce the range of dynamical mechanisms that give rise to metastable dynamics. Not all forms of metastability are possible with this assumption. Some forms of metastability, such as those associated with chaotic itineracy, require significantly more complex attractor structures to emerge.

We justify this assumption in our work for two primary reasons. Firstly, in the models we have used, the individual attractor states that represent the behaviors may be substituted with more complex attractors as in [56]. Secondly, work in topological dynamics shows that although state transitions may be caused by a range of structures, the basic building blocks for such transitions are due to proximity to a bifurcation or the formation of heteroclinic-like structural symmetries [15]. This means that although our analysis is far from exhaustive, the insights gleamed will have implications for most dynamical systems with a metastable structure. However, a fruitful direction of future work could extend this work to more complex state structures.

In applying these methods to experimental systems, we are invariably faced with noise effects. This makes it difficult to distinguish between which mechanisms may be responsible for behavior switching in these systems. Some mathematical and computational progress is needed to develop techniques for distinguishing between these mechanisms under noisy conditions. However, as we and others [49] have shown, the robustness to noise itself may be an indicator of the underlying dynamical mechanisms. Additional work has also found that synchronization with oscillatory inputs may also be an important means of distinguishing among the various mechanisms [49].

Finally, we turn to the implications of this work for our theoretical understanding of adaptive behavior. There is a growing appreciation that behavior is shaped by interactions of processes occurring at a variety of scales [57]. This forces us to re-examine how we conceive of behavioral switching, not as the sequencing of distinct modes triggered by external stimuli, but as temporary synergies of coordinated activity unfolding within an ongoing physiological process. In *C. elegans*, for example, forward-to-reversal switching may be better understood not as a response to a change in stimulus, but as an expression of its intrinsic physiological cycles.

The mathematical machinery we have presented provides a way to formalize this shift in perspective. By treating behavioral states as metastable rather than multistable, we model switching as arising from the internal dynamics of the organism rather than from transitions between externally defined stimulus–response states. In this way, dynamical systems theory offers a framework for integrating physiological cycles with temporary behavioral synergies. This approach allows us to understand behavior and physiology not as separable domains, but as mutual constraints on a single multiscale dynamical process.

## 8 Acknowledgments

We would like to thank Lindsay Stolting, Connor McShaffrey, and Luis Favela for their helpful comments on the paper, and Quan Le Thien for his suggestions on plotting. We also thank the Computational Neuroethology Lab at Indiana University for helpful discussions during the development of this paper.

## 9 Appendix

### 9.1 CTRNN Construction Method

In [40], the authors demonstrate how a transition graph can be embedded as a network of solutions to a set of differential equations through the use of only a few parameters. They first prove this is true when the standard sigmoid nonlinearity is replaced by a three-part piecewise smooth function:

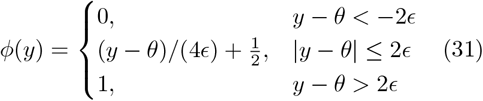

The authors show that setting the weight matrix as below will embed the graph into the dynamics:

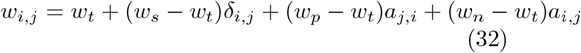

where *w*_*s*_, *w*_*t*_, *w*_*p*_, *w*_*n*_ are the network parameters. In this work, we do not use lateral states and thus we set *w*_*t*_ = 0. The function *δ*_*i,j*_ returns some positive value (which by abuse of, we refer to as *δ*) if there is a connection from *i, j* in the graph that is being embedded, and 0 otherwise.

The conditions necessary for the parameters can be state as:

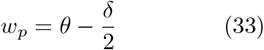

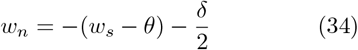

The variable *θ* is the bias that is shared across all nodes of the CTRNN which in our case is 0. Under these conditions, variations of the parameter *w*_*p*_ (or *w*_*n*_) will posses a saddle node bifurcation which forms a multistable network. The authors conjecture that the other side of the bifurcation exhibits a limit cycle.

This approach cannot account for self-loops (a node with an edge to itself) or two-loops (two cells with edges to one another) because it uses a similar logic to the simplex method. In both cases, the switching dynamics rely on inhibition from successor cells to the predecessor cell, which is incompatible with the occurrence one- and two-loops. This method also does not support Δ-cliques where two cells have a direct connection as well as a one-step indirect connection in the same direction because they result in state skipping in the dynamics.

The authors then demonstrate that this method also works when the function *f* is the standard sigmoid nonlinearity. In the case of two nodes, they explicitly work out the location of the saddle node bifurcation for a given set of gains and biases. They numerically demonstrate similar results for several larger networks for 2, 3, 4, and 10 cell CTRNNs.

#### 9.2 GLV Dwell-Time Equations

In [46], the authors develop a reparameterization of the GLV system that is amenable to explicit control of the dwell time of a given state. In this section, we explain this reparameterization and show how it can be used to control dwell times as we did in equation (18).

We start with the GLV system with the added growth parameter:

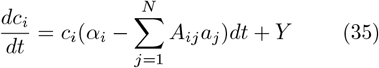

If we define *A* such that:

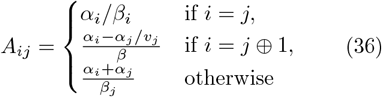

We can explicitly solve for the eigenvalue of the manifold going from cell *i* to cell *j* as:

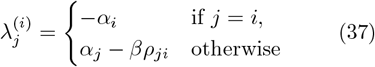

This is useful because, we can frame the dwell time of a trajectory near a node on the heteroclinic network as the mean first passage time from the stable manifold of that node to the stable manifold of the next node. An integral expression for this mean first passage time was defined by Stone and Holmes in [58]:

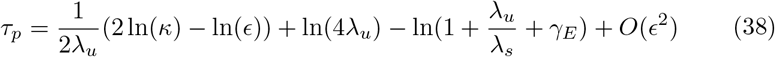

In this equation the mean first passage time *τ*_*p*_ is expressed as a function of the eigenvalues of the unstable manifold of the dwelling node, *λ*_*u*_, the eigenvalue of the stable manifold of the next node *λ*_*s*_, the variance of the noise term *ϵ*, some chosen parameter *κ* and the Euler-Mascheroni constant, *γ*_*E*_. *κ* for this work we choose *κ* = 10^−1^ as suggested in [46].

Finally, we can plug in the equations for the eigenvalues as in equation (21) to expand and solve for the desired dwell time *τ* :

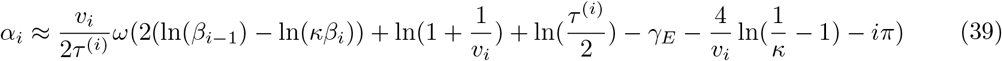

Choosing *β*_*i*_ = 1 and *v*_*i*_ = 1, we can explicitly solve for the value of *α*_*i*_ such that the matrix *A* produces. the desired dwell times.

## References

[1] Mora, T. & Bialek, W. Are biological systems poised at criticality? Journal of Statistical Physics 144, 268–302 (2011). URL http://arxiv.org/abs/1012.2242. 1012.2242 [q-bio].

[2] Hogan, N. & Sternad, D. Dynamic primitives of motor behavior. Biological Cybernetics 106, 727–739 (2012). URL http://link.springer.com/10.1007/s00422-012-0527-1.

[3] Mathis, A. et al. DeepLabCut: markerless pose estimation of user-defined body parts with deep learning. Nature Neuroscience 21, 1281–1289 (2018). URL https://www.nature.com/articles/s41593-018-0209-y.

[4] Pereira, T. D. et al. SLEAP: A deep learning system for multi-animal pose tracking. Nature Methods 19, 486–495 (2022). URL https://www.nature.com/articles/s41592-022-01426-1. Publisher: Nature Publishing Group.

[5] Datta, S. R., Anderson, D. J., Branson, K., Perona, P. & Leifer, A. Computational Neuroethology: A Call to Action. Neuron 104, 11–24 (2019). URL https://linkinghub.elsevier.com/retrieve/pii/S0896627319308414.

[6] Kato, S. et al. Global Brain Dynamics Embed the Motor Command Sequence of Caenorhabditis elegans. Cell 163, 656– 669 (2015). URL https://linkinghub.elsevier.com/retrieve/pii/S0092867415011964.

[7] Costa, A. C., Ahamed, T., Jordan, D. & Stephens, G. J. A Markovian dynamics for Caenorhabditis elegans behavior across scales. Proceedings of the National Academy of Sciences 121, e2318805121 (2024). URL https://www.pnas.org/doi/full/10.1073/pnas.2318805121. Publisher: Proceedings of the National Academy of Sciences.

[8] Roberts, W. M. et al. A stochastic neuronal model predicts random search behaviors at multiple spatial scales in C. elegans. eLife 5, e12572 (2016). URL 10.7554/eLife.12572. Publisher: eLife Sciences Publications, Ltd.

[9] Costa, A. C., Sridhar, G., Wyart, C. & Vergassola, M. Fluctuating Landscapes and Heavy Tails in Animal Behavior. PRX Life 2, 023001 (2024). URL https://link.aps.org/doi/10.1103/PRXLife.2.023001. Publisher: American Physical Society.

[10] Bialek, W. On the dimensionality of behavior. Proceedings of the National Academy of Sciences 119, e2021860119 (2022). URL https://www.pnas.org/doi/10.1073/pnas.2021860119. Publisher: Proceedings of the National Academy of Sciences.

[11] Chiel, H. J. & Beer, R. D. The brain has a body: adaptive behavior emerges from interactions of nervous system, body and environment. Trends in Neurosciences 20, 553–557 (1997). URL https://www.cell.com/trends/neurosciences/abstract/S0166-2236(97)01149-1.

[12] Hancock, F. et al. Metastability demystified — the foundational past, the pragmatic present and the promising future. Nature Reviews Neuroscience 26, 82–100 (2025). URL https://www.nature.com/articles/s41583-024-00883-1.

[13] Koch, D., Nandan, A., Ramesan, G. & Koseska, A. Biological computations: Limitations of attractor-based formalisms and the need for transients. Biochemical and Biophysical Research Communications 720, 150069 (2024). URL https://www.sciencedirect.com/science/article/pii/S0006291X24006053.

[14] Rossi, K. L. et al. A unified framework of metastability in neuroscience (2023). URL http://arxiv.org/abs/2305.05328. 2305.05328 [nlin, q-bio].

[15] Gorban, A. N. Singularities of Transition Processes in Dynamical Systems: Qualitative Theory of Critical Delays (2004). URL http://arxiv.org/abs/chao-dyn/9703010. chao-dyn/9703010.

[16] Atanas, A. A. et al. Brain-wide representations of behavior spanning multiple timescales and states in C. elegans. Cell 186, 4134–4151.e31 (2023). URL https://www.cell.com/cell/abstract/S0092-8674(23)00850-4. Publisher: Elsevier.

[17] Schlegel, P. et al. Whole-brain annotation and multi-connectome cell typing of Drosophila. Nature 634, 139–152 (2024). URL https://www.nature.com/articles/s41586-024-07686-5. Publisher: Nature Publishing Group.

[18] Stephens, G. J., Johnson-Kerner, B., Bialek, W. & Ryu, W. S. Dimensionality and Dynamics in the Behavior of C. elegans. PLOS Computational Biology 4, e1000028 (2008). URL https://journals.plos.org/ploscompbiol/article?id=10.1371/journal.pcbi.1000028. Publisher: Public Library of Science.

[19] Cohen, N. & Denham, J. E. Whole animal modeling: piecing together nematode locomotion. Current Opinion in Systems Biology 13, 150–160 (2019). URL https://www.sciencedirect.com/science/article/pii/S2452310018301082.

[20] Kim, J., Santos, J. A., Alkema, M. J. & Shlizerman, E. Whole integration of neural connectomics, dynamics and biomechanics for identification of behavioral sensorimotor pathways in Caenorhabditis elegans (2019). URL https://www.biorxiv.org/content/10.1101/724328v1. Pages: 724328 Section: New Results.

[21] Ahamed, T., Costa, A. C. & Stephens, G. J. Capturing the continuous complexity of behaviour in Caenorhabditis elegans. Nature Physics 17, 275–283 (2021). URL https://www.nature.com/articles/s41567-020-01036-8.

[22] Izquierdo, E. J. & Beer, R. D. The whole worm: brain-body-environment models of C. elegans. Current Opinion in Neurobiology 40, 23–30 (2016).

[23] Dahlberg, B. A. & Izquierdo, E. J. Contributions from parallel strategies for spatial orientation in C. elegans (2020). URL https://direct.mit.edu/isal/proceedings-abstract/isal2020/32/16/98418.

[24] Morrison, M. & Young, L.-S. A data-driven biophysical network model reproduces C. elegans premotor neural dynamics (2024). URL http://arxiv.org/abs/2501.00278. 2501.00278 [q-bio].

[25] Olivares, E., Izquierdo, E. J. & Beer, R. D. A neuromechanical model of multiple network oscillators for forward locomotion in C. elegans. bioRxiv 710566 (2019). URL https://www.biorxiv.org/content/10.1101/710566v2. Publisher: Cold Spring Harbor Laboratory Section: New Results.

[26] Morrison, M. & Young, L.-S. Chaotic heteroclinic networks as models of switching behavior in biological systems. Chaos: An Interdisciplinary Journal of Nonlinear Science 32, 123102 (2022). URL http://arxiv.org/abs/2208.10654. 2208.10654 [nlin].

[27] Ashwin, P., Coombes, S. & Nicks, R. Mathematical Frameworks for Oscillatory Network Dynamics in Neuroscience. The Journal of Mathematical Neuroscience 6, 2 (2016). URL 10.1186/s13408-015-0033-6.

[28] Ashwin, P., Fadera, M. & Postlethwaite, C. Network attractors and nonlinear dynamics of neural computation. Current Opinion in Neurobiology 84, 102818 (2024). URL https://www.sciencedirect.com/science/article/pii/S0959438823001435.

[29] Rabinovich, M. I., Huerta, R., Varona, P. & Afraimovich, V. S. Transient Cognitive Dynamics, Metastability, and Decision Making. PLOS Computational Biology 4, e1000072 (2008). URL https://journals.plos.org/ploscompbiol/article?id=10.1371/journal.pcbi.1000072. Publisher: Public Library of Science.

[30] Beer, R. D. & Gallagher, J. C. Evolving Dynamical Neural Networks for Adaptive Behavior. Adaptive Behavior 1, 91– 122 (1992). URL 10.1177/105971239200100105. Publisher: SAGE Publications Ltd STM.

[31] Lin, S. et al. Characterizing the structure of mouse behavior using Motion Sequencing. Nature Protocols 19, 3242–3291 (2024). URL https://www.nature.com/articles/s41596-024-01015-w.

[32] Beer, R. D. Parameter Space Structure of Continuous-Time Recurrent Neural Networks. Neural Computation 18, 3009–3051 (2006). URL https://direct.mit.edu/neco/article/18/12/3009-3051/7104.

[33] Castro, S.B.S.D. & Lohse, A. Construction of heteroclinic networks in \mathbbR4. Nonlinearity 29, 3677 (2016). URL 10.1088/0951-7715/29/12/3677. Publisher: IOP Publishing.

[34] Postlethwaite, C. & Ashwin, P. On designing heteroclinic networks from graphs. Physica D: Nonlinear Phenomena 265, 26–39 (2013). URL https://www.sciencedirect.com/science/article/abs/pii/S0167278913002601. Publisher: North-Holland.

[35] Afraimovich, V. S., Rabinovich, M. I. & Varona, P. Heteroclinic Contours in Neural Ensembles and the Winnerless Competition Principle. International Journal of Bifurcation and Chaos 14, 1195–1208 (2004). URL http://arxiv.org/abs/nlin/0304016. nlin/0304016.

[36] Guckenheimer, J. & Holmes, P. Nonlinear Oscillations, Dynamical Systems, and Bifurcations of Vector Fields Applied Mathematical Sciences (Springer, New York, NY, 1983). URL http://link.springer.com/10.1007/978-1-4612-1140-2. ISSN: 0066-5452, 2196-968X.

[37] Rabinovich, M., Bick, C. & Varona, P. Beyond neurons and spikes: cognon, the hierarchical dynamical unit of thought. Cognitive Neurodynamics 18, 3327–3335 (2024). URL 10.1007/s11571-023-09987-3.

[38] Komarov, M. A., Osipov, G. V. & Zhou, C. S. Heteroclinic contours in oscillatory ensembles. Physical Review E 87, 022909 (2013). URL https://link.aps.org/doi/10.1103/PhysRevE.87.022909. Publisher: American Physical Society.

[39] Meyer-Ortmanns, H. Heteroclinic networks for brain dynamics. Frontiers in Network Physiology 3, 1276401 (2023). URL https://www.ncbi.nlm.nih.gov/pmc/articles/PMC10663269/.

[40] Ashwin, P. & Postlethwaite, C. Excitable networks for finite state computation with continuous time recurrent neural networks. Biological Cybernetics 115, 519–538 (2021). URL https://www.ncbi.nlm.nih.gov/pmc/articles/PMC8589808/.

[41] Chow, C. C. & Karimipanah, Y. Before and beyond the Wilson–Cowan equations. Journal of Neurophysiology 123, 1645–1656 (2020). URL https://journals.physiology.org/doi/full/10.1152/jn.00404.2019. Publisher: American Physiological Society.

[42] Beer, R. D. The Global Structure of Codimension-2 Local Bifurcations in Continuous-Time Recurrent Neural Networks. Biological Cybernetics 116, 501– 515 (2022). URL http://arxiv.org/abs/2111.04547. 2111.04547 [q-bio].

[43] Beer, R. D.Barwich, A.-S. & Severino, G. J. Milking a spherical cow: Toy models in neuroscience. The European Journal of Neuroscience 60, 6359–6374 (2024).

[44] Rackauckas, C. & Nie, Q. DifferentialEquations.jl – A Performant and Feature-Rich Ecosystem for Solving Differential Equations in Julia. Journal of Open Research Software 5 (2017). URL https://openresearchsoftware.metajnl.com/articles/10.5334/jors.151.

[45] Armbruster, D., Stone, E. & Kirk, V. Noisy heteroclinic networks. Chaos: An Interdisciplinary Journal of Nonlinear Science 13, 71–79 (2003). URL https://pubs.aip.org/cha/article/13/1/71/511017/Noisy-heteroclinic-networks.

[46] Horchler, A. D., Daltorio, K. A., Chiel, H. J. & Quinn, R. D. Designing responsive pattern generators: stable heteroclinic channel cycles for modeling and control. Bioinspiration & Biomimetics 10, 026001 (2015). URL 10.1088/1748-3190/10/2/026001. Publisher: IOP Publishing.

[47] Kirk, V. & Silber, M. A competition between heteroclinic cycles. Nonlinearity 7, 1605–1621 (1994). URL http://www.scopus.com/inward/record.url?scp=22244472608&partnerID=8YFLogxK.

[48] Maisel, B. Winnerless Competition Principle in Individual Neurons and Cognitive Networks.

[49] Koch, D. & Koseska, A. Ghost cycles exhibit increased entrainment and richer dynamics in response to external forcing compared to slow-fast systems (2024). URL http://arxiv.org/abs/2403.19624. 2403.19624 [nlin].

[50] Beer, R. D. Dynamica: A Mathematica package for the analysis of smooth dynamical systems. https://rdbeer.pages.iu.edu (2021). Version 1.0.11, xreleased May 8, 2021.

[51] Jaramillo, G., Mrad, L. & Stepien, T. L. Dynamics of a linearly-perturbed May-Leonard competition model. Chaos: An Interdisciplinary Journal of Nonlinear Science 33, 063121 (2023). URL http://arxiv.org/abs/2210.04342. 2210.04342 [math].

[52] Phillipson, P. E., Schuster, P. & Johnston, R. G. An Analytic Study of the May-Leonard Equations. SIAM Journal on Applied Mathematics 45, 541–554 (1985). URL http://epubs.siam.org/doi/10.1137/0145032.

[53] Nechyporenko, K., Ashwin, P. & Tsaneva-Atanasova, K. A Novel Route to Oscillations via non-central SNICeroclinic Bifurcation: unfolding the separatrix loop between a saddle-node and a saddle (2024). URL http://arxiv.org/abs/2412.12298. 2412.12298 [math].

[54] Rouse, N. A., Horchler, A. D., Chiel, H. J. & Daltorio, K. A. Stable heteroclinic channels as a decision-making model: overcoming low signal-to-noise ratio with mutual inhibition. Bioinspiration & Biomimetics 20, 036004 (2025). URL https://iopscience.iop.org/article/10.1088/1748-3190/adc057/meta. Publisher: IOP Publishing.

[55] Hansen, N. The CMA Evolution Strategy: A Tutorial (2023). URL http://arxiv.org/abs/1604.00772. 1604.00772 [cs].

[56] Dellnitz, M., Field, M., Golubit-Sky, M., Ma, J. & Hohmann, A. CYCLING CHAOS. International Journal of Bifurcation and Chaos (2011). URL https://www.worldscientific.com/worldscinet/ijbc. Publisher: World Scientific Publishing Company.

[57] Favela, L. H. The Ecological Brain: Unifying the Sciences of Brain, Body, and Environment (Routledge, New York, 2023).

[58] Stone, E. & Holmes, P. Random Perturbations of Heteroclinic Attractors. SIAM Journal on Applied Mathematics (2006). URL https://epubs.siam.org/doi/10.1137/0150043.

